# Phylomedicine of mutational processes in somatic cancer cell populations

**DOI:** 10.1101/2021.04.02.438268

**Authors:** Sayaka Miura, Tracy Vu, Jiyeong Choi, Sudhir Kumar

## Abstract

Mutational processes in somatic cancer cell populations are constantly changing, leaving their signatures in the accumulated genomic variation in tumors. The inference of mutational signatures from the observed genetic variation enables spatiotemporal tracking of tumor mutational processes that evolve due to cellular environmental changes, mutations, and treatment regimes. Ultimately, mutational patterns illuminate the mechanistic understanding of their evolution in cancer progression. We show that the integration of cancer cell phylogeny with mutational signature deconvolution enables higher-resolution detection of gain and loss of mutational processes within the phylogeny. This approach to analyzing somatic genomic variations in 61 lung cancer patients revealed a high turn-over of mutational processes over time and closely related clonal lineages. Some mutational signatures (e.g., smoking-related) showed a higher propensity to be lost, whereas others (e.g., AID/APOBEC) were gained during lung tumors evolution. These observations shed light on the evolution of mutational processes in somatic cell evolution.

## INTRODUCTION

Tumor cells accumulate somatic mutations during cancer progression, in which cells exhibit dynamic demography, including emergence, expansion, and extinction (Bailey et al., 2018; Martincorena & Campbell, 2015). Through the analysis of genomic variation, researchers now routinely reconstruct mutational histories and phylogenies of clones (Brown et al., 2017; El-Kebir et al., 2018; Miura et al., 2020; Turajlic et al., 2018; Zhao et al., 2016). In a clone phylogeny, variants can be localized to individual branches and relative frequencies of different variant types compared across lineages to detect shifts in cellular mutational processes over time. For example, the trunk of the clone phylogeny in **figure 1** shows many more C→A transversions than in its descendants, suggesting that the process of mutagenesis is not the same over time in this lung cancer patient. Such temporal comparisons of genome variation patterns are a new frontier in enhanced understanding of the intricacies of evolution in individual tumors and patients. These comparisons reveal how pre-existing genetic alterations and treatment regimens are fundamentally altering the landscape of mutational processes, often producing resistant cells that promote cancer progression (Ashley et al., 2019; Barry et al., 2018; de Bruin et al., 2014; Dentro et al., 2020; Gerstung et al., 2020; Leong et al., 2019).

**Figure 1.**
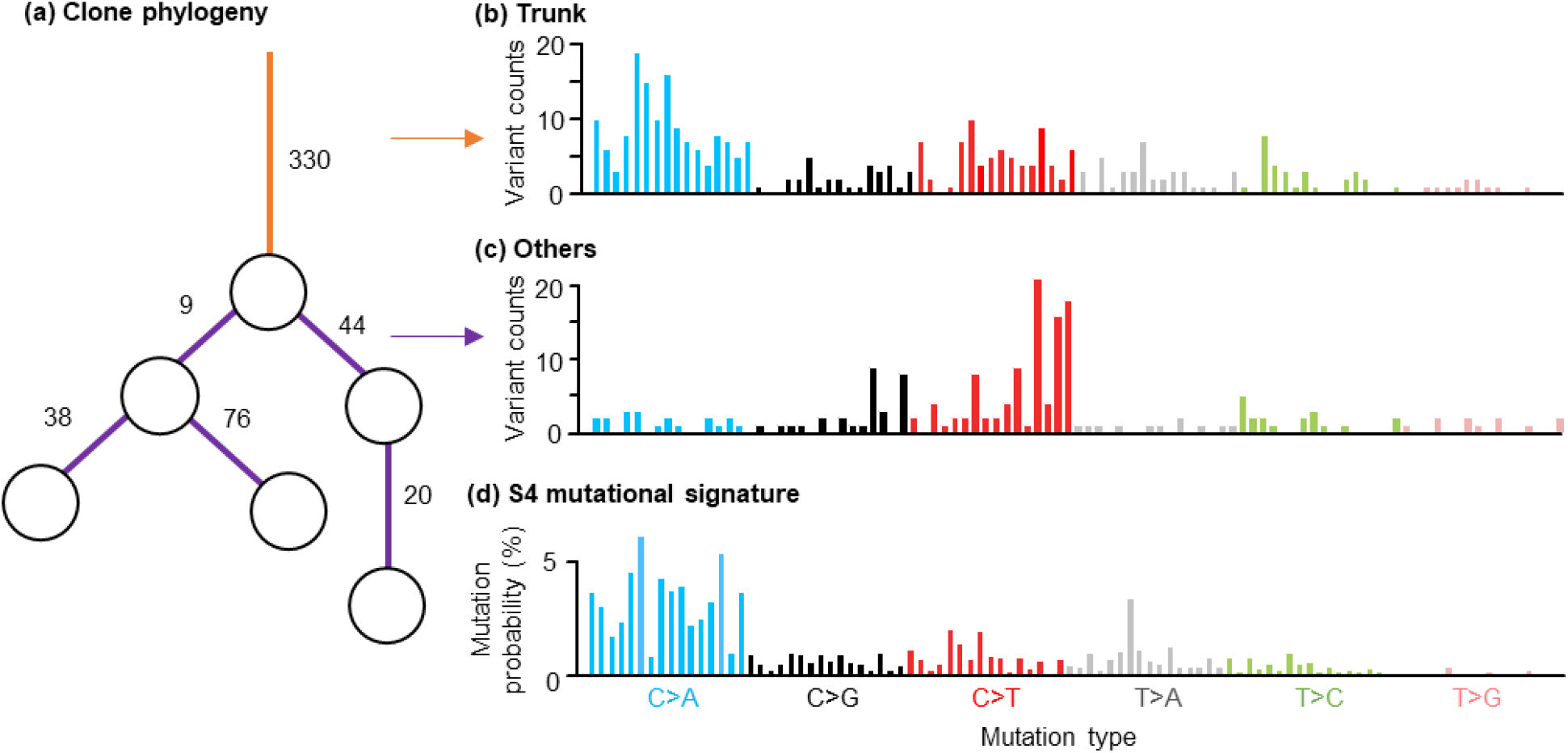
Clone phylogeny and variant counts from lung cancer data. (**a**) Clone phylogeny of 6 clones. Clones are shown with circles. Numbers along branches represent variant counts. (**b** and **c**) Observed variant counts of the trunk (orange branch; **b**) and the other branches (purple; **c**). The data was obtained from Jamal-Hanjani et al. (2017) (CRUK0025 dataset). (**d**) COSMIC signature S4, which is characterized by many C to A mutations.

Many mutational processes leave distinct signatures in the form of types of variants and their relative counts. For example, a large C→A variant frequency is a tell-tale sign of smoking-related mutational processes that arise early (COSMIC signature S4; **Fig. 1b** and **1d**). Their activity begins to decline after smoking cessation (Alexandrov et al., 2016; Le Calvez et al., 2005). In contrast, age-related mutagenic processes create C→T transitions that arise throughout a person’s lifetime (COSMIC signature S1) and result in the decay of methylated CpG sites (Alexandrov et al., 2013; Alexandrov et al., 2018; Alexandrov & Stratton, 2014; Van Hoeck et al., 2019). Many distinct mutational signatures have been inferred from the genetic variation found in various cancers’ tumors, which has been assembled in online catalogs (Alexandrov et al., 2020; Goncearenco et al., 2017). For example, 30 signatures have been recognized in COSMIC v2, each of which is a vector of 96 different mutational contexts consisting of the mutated base and adjacent 5’ and 3’ bases (e.g., **Fig. 1d**) (Alexandrov et al., 2015; Alexandrov et al., 2020; Tate et al., 2019).

Computational methods are available to estimate their relative activities of mutational signatures from observed tumor genetic variants (Blokzijl et al., 2018; Huang et al., 2018; Rosenthal et al., 2016). Mutational processes operating in early and late stages of cancer progression have been contrasted using predicted mutational signatures (Ashley et al., 2019; Barry et al., 2018; de Bruin et al., 2014; Dentro et al., 2020; Gerstung et al., 2020; Leong et al., 2019). Researchers have also begun to analyze branch-specific mutational signatures in clone phylogenies to discover mutagens linked with the origin of new clones in cancer patients (Barry et al., 2018; Hao et al., 2016; Roper et al., 2019; Wang et al., 2019).

Successful mutational signature detection using existing methods currently requires hundreds of somatic variants (Li et al., 2018). This requirement makes the inference of evolutionary dynamics of mutational signatures at a finer phylogenetic resolution (e.g., branch-by-branch) challenging because the collection of variants in individual branches in clone phylogenies is often small (**Fig. 2** and **3**). For example, fewer than 100 variants were mapped to most branches in carcinoma cell phylogenies (Jamal-Hanjani et al., 2017) (**Fig. 2** and **3**). It is not yet possible for such small collections of variants to detect branch-specific mutational signatures reliably (Li et al., 2018). This problem is illustrated in an analysis of a simulated clone phylogeny modeled after an adenocarcinoma clone phylogeny (**Fig. 4a**; phylogeny CRU0079 in **Fig. 2**). The available state-of-the-art methods used to branch-specific variants frequently produced too many signatures, while some correct signatures remained undetected (**Fig. 4b-d**). This means that we cannot yet reliably detect the evolution of branch-specific signatures over time in a patient, limiting us to gross comparisons that pool variants to build large-enough collections (de Bruin et al., 2014; Dong et al., 2018; Hao et al., 2016; Nahar et al., 2018).

**Figure 2.**
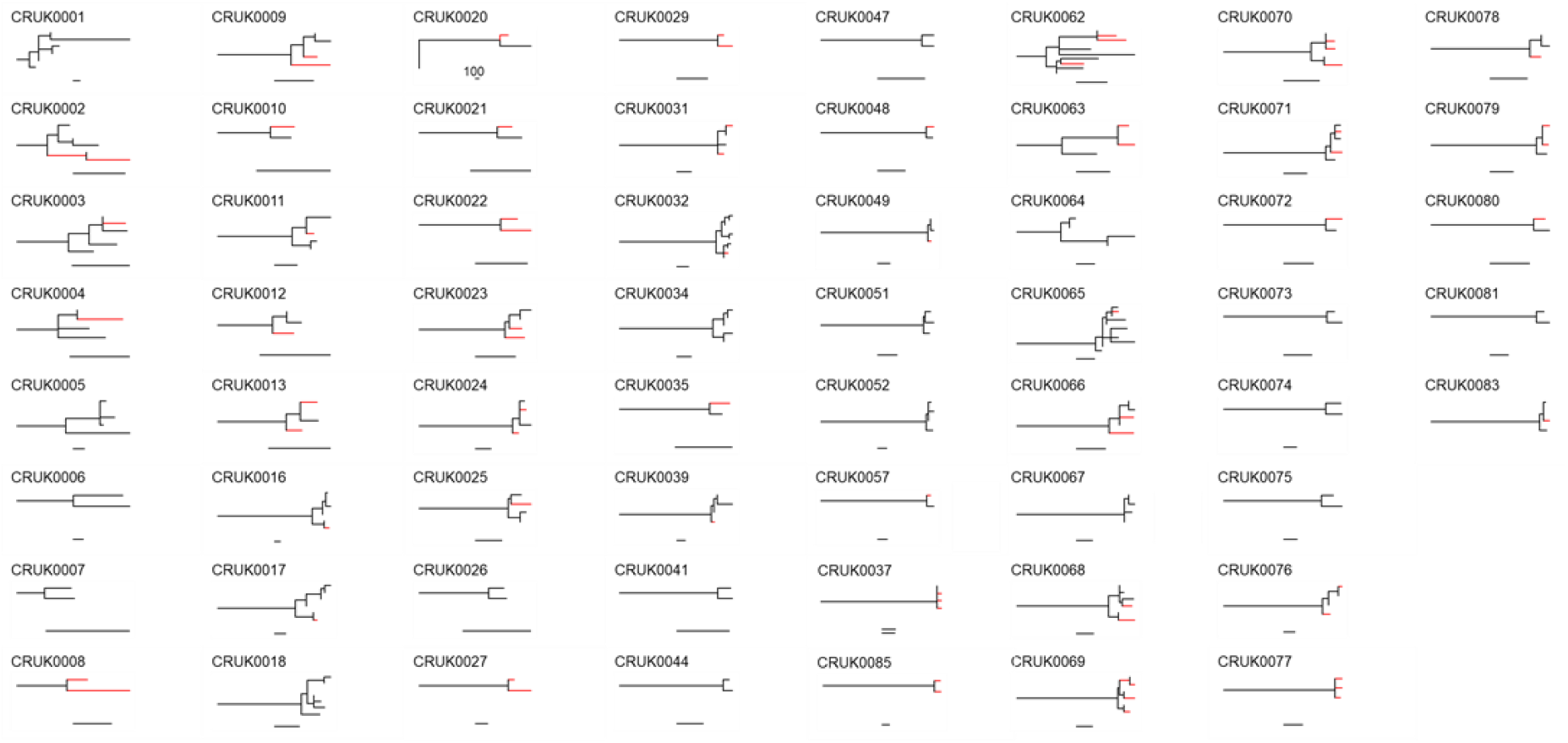
Clone phylogenies from data in Jamal-Hanjani et al. (2017). Only phylogenies with >1 tip were included. Branches with <20 variants were combined for signature detection (see **Methods**). Combined branches were shown with red. The scale bar is equal to 100 variants in each case.

**Figure 3.**
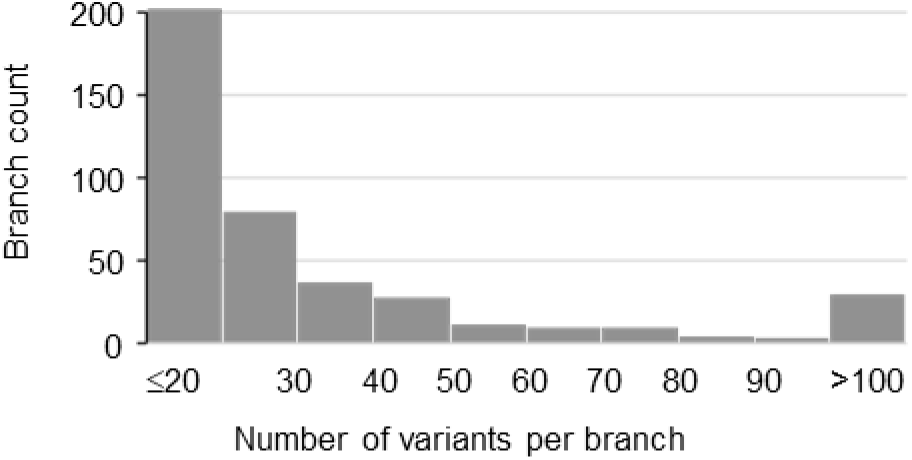
The number of variants in individual branches of all the clone phylogenies that are shown in Figure 2. Clone phylogenies were obtained from data in Jamal-Hanjani et al. (2017).

**Figure 4.**
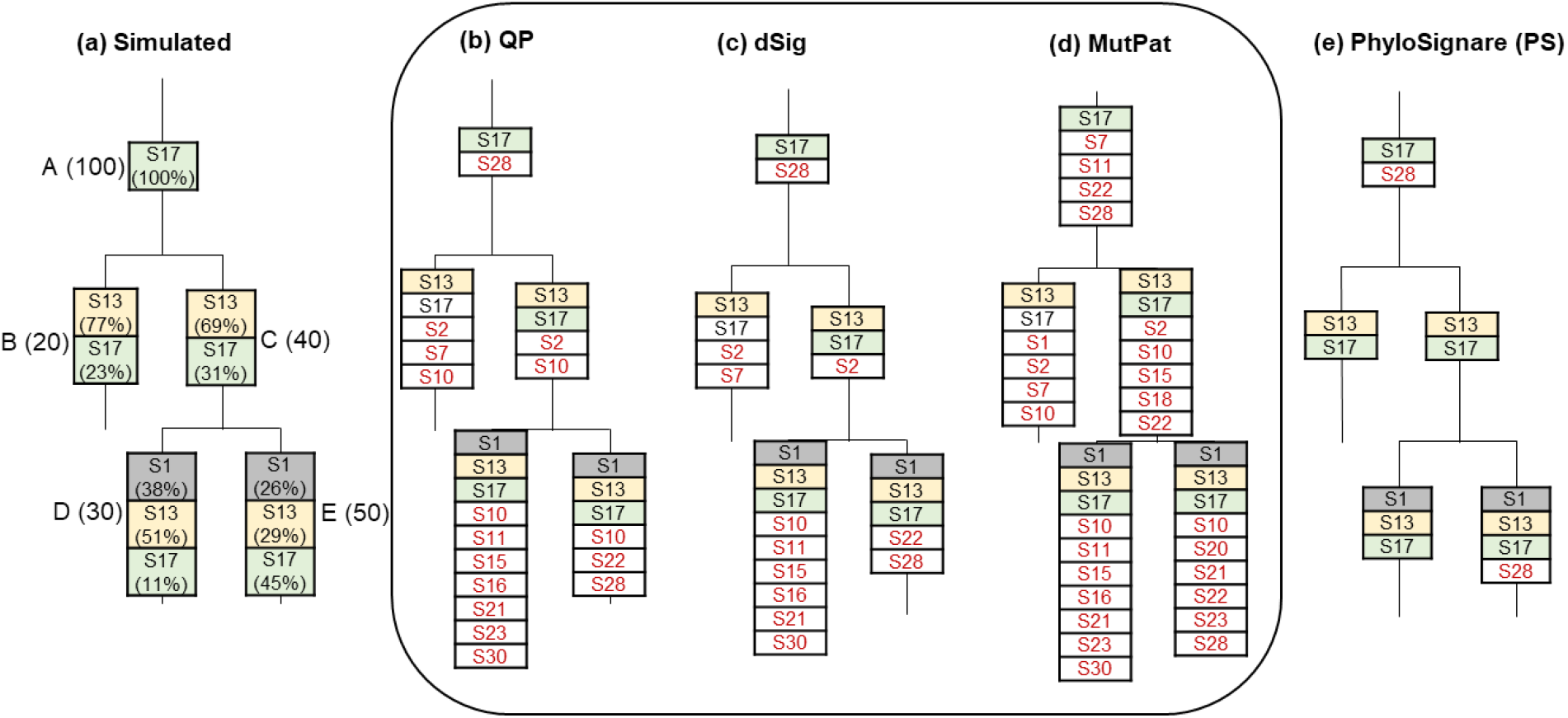
Mutational signatures detected by different methods for individual branches in the simulated clone phylogeny. (**a**) Model clone phylogeny and simulated mutational signatures. There are five branches: A – E with 20 – 100 variants (counts in parentheses next to the branch name) and each signature’s relative activity (shown below the signature name). (**b-d**) mutational signatures inferred by using different methods: (**b**) QP, (**c**) dSig, and (**d**) MutPat. (**e**) Mutational signatures inferred by applying the *PhyloSignare* approach with the QP Method. Incorrectly detected signatures are shown with red letters, and correct signatures not detected are shown in a white box with black letters.

We hypothesized that the detection of mutational signatures would be more accurate if the clone phylogeny is utilized alongside mutation signature detection approaches. This idea is based on the expectation that neighboring branches in the clone phylogeny will share some mutational signatures due to their shared environment and evolutionary history, e.g., Dentro et al. (2018). We leveraged this property to infer branch-specific mutational signatures through a joint analysis of the collection of mutations mapped on phylogenetically proximal branches of the clone phylogeny, which is called *PhyloSignare* and presented below. Then, we present an assessment of *PhyloSignare*’s usefulness by analyzing computer-simulated datasets, which establish that *PhyloSignare* can significantly improve the accuracy of current methods for smaller collections of variants (Blokzijl et al., 2018; Huang et al., 2018; Rosenthal et al., 2016). Finally, we apply *PhyloSignare* to infer mutational signature evolution in non-small cell lung cancer patients, revealing branch-specific mutational signatures at a finer phylogenetic resolution.

## RESULTS

### The *PhyloSignare* (PS) approach

As noted above, current methods produce many spurious signatures when the number of variants analyzed is not large enough. To detect spurious signatures, we estimate an importance score (iS) for each signature predicted using an existing method, e.g., a quadratic programming approach (QP) (Huang et al., 2018). This score contrasts the statistical fit of the predicted signatures to explain the frequencies of branch-specific variants with and without the given signature (see *Methods* section for details). When iS is small, the predicted signature may be spurious. For example, iS2, iS7, and iS10 for signatures S2, S7, and S10, respectively, were very small (< 0.02) in branch B for the computer-simulated dataset. None of these signatures were present in this branch (**Fig. 4a**). In contrast, the correct signature S13 received a high score (iS13 = 0.87). Therefore, iS can detect spurious candidate signatures in branch-by-branch analysis, potentially reducing the false-positive detection rate.

However, the above procedure does not recover signature S17 in branch B analysis (**Fig. 4a**). To reduce such false negatives, we pool variants from neighboring branches in the clone phylogeny to increase the data size, as current mutational signature methods work better with a larger number of variants. For example, pooling variants in branch B with its ancestral branch (trunk, branch A) expands the variant collection to 120 variants. Now QP predicts the presence of S17 along with a few false positives (S7, S10, and S28). This process is repeated by pooling variants from other branch B neighbors, and then a candidate list of signatures is generated for branch B (S2, S7, S10, S13, S17, and S28). After that, we use the iS approach to identify potentially spurious signatures and exclude them. This process is applied to every branch in the clone phylogeny to infer a candidate list of signatures.

In the final step, we estimate the relative activities of signatures in every branch. For a given branch, we consider all signatures inferred for that branch (as above) as well as the signatures of its immediate relatives (ancestor and sibling branches). Then, QP is used to estimate activities for all candidate signatures in each branch. We found this pooling of candidate signatures to minimize spurious gain and loss of signatures caused by small sample sizes. The estimated activity of incorrect signatures was usually zero or nearly so (i.e., the false-positive rate did not increase), and additional correct signatures were found, reducing the false-negative rate (see *Methods*). In the current example, *PhyloSignare* improved the accuracy of mutational signatures detected by the QP method for every branch (PS_QP_; **Fig. 4e**). Beyond the illustrative example, we evaluated the benefit of using *PhyloSignare* for many more datasets produced using computer simulation and three signature detection methods (QP, dSig, and MutPat) to obtain a more general trend.

### Accuracy of PhyloSignare approach

We tested the performance of *PhyloSignare* by analyzing 1,080 branches in 180 clone phylogenies that were simulated and utilized by others (see *Methods*). Clone phylogenies consisted of five or seven branches, with fewer than 100 variants mapped to 486 branches (out of 1,080). In these simulations, signatures were randomly sampled from 30 COSMIC signatures (v2), and up to two branches experienced loss and/or gain of as many as eight signatures (Christensen et al., 2020). On average, the precision of *PhyloSignare* was 93%, a large improvement over the direct use of QP (66%; **Fig. 5a**). *PhyloSignare* coupled with the dSig method (Rosenthal et al., 2016) was also more accurate (92% vs. 70%). A similar performance gain (92% vs. 65%) was seen when using MutPat (Blokzijl et al., 2018). This gain in precision did not compromise recall significantly (**Fig. 5b**). Consequently, overall accuracy (F1) was much better when *PhyloSignare* was coupled with signature detection methods (**Fig. 5c**). As expected, overall accuracy (F1) declined with the decreasing number of variants (**Fig. 5f**), but *PhyloSignare* showed more than 80% accuracy even for short branches. The true positive rate (precision) of *PhyloSignare* was very similar across variant sample sizes (**Fig. 5d**), but the recall was negatively impacted by lower sample sizes (**Fig. 5e**).

**Figure 5:**
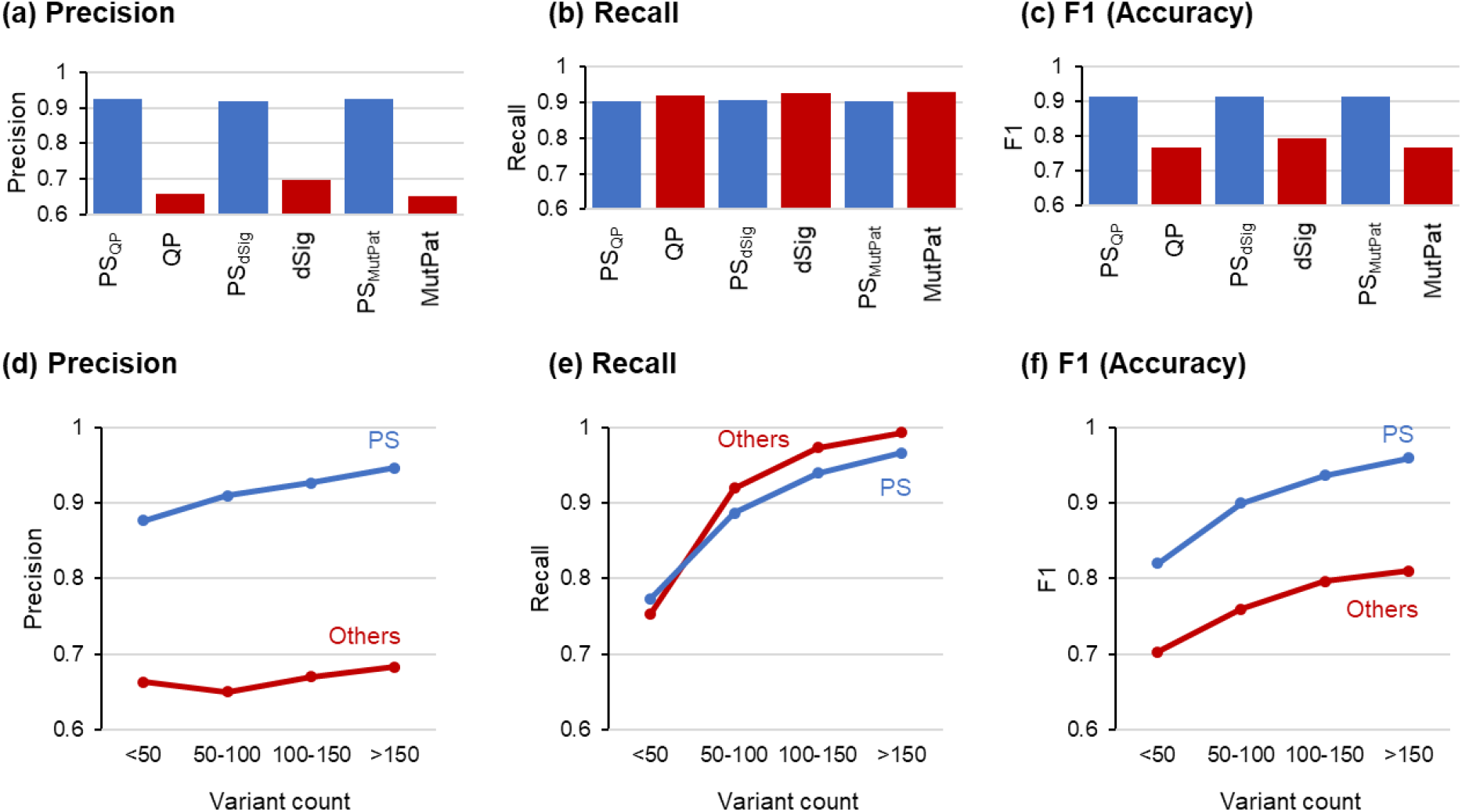
The performance of *PhyloSignare*. (**a**) Precision, (**b**) recall, and (**c**) F1 score for all the signatures across all datasets for QP, dSig, and MutPat without and with *PhyloSignare* (PS) approach (PS_QP_, PS_dSig_, and PS_MutPat_, respectively). (**d)** Precision, (**e**) recall, and (**f**) F1 scores for all the signatures for branches of various lengths. Signatures were pooled across all datasets in the computation. Precision was computed as the number of correct signatures detected divided by the total number of signatures detected. The recall was the number of correct signatures detected divided by the total number of simulated signatures. F1 = 2× Precision×Recall/(Precision+Recall).

### Dynamics of mutational signatures in lung cancer patients

We next analyzed 61 clone phylogenies (see *Methods*) by using *PhyloSignare*. In these phylogenies, the number of variants was generally less than 100 for individual branches (**Fig. 2** and **3**). We begin by describing results from the analysis of one lung adenocarcinoma patient (CRUK0025; **Fig. 6**). This patient’s clone phylogeny consisted of six branches, with branch A (trunk) containing 330 variants and fewer than 100 variants mapped to all other branches. In the trunk, *PhyloSignare* predicted the presence of S4, a signature of a smoking-related mutational process that produces many C→A variants (**Fig. 6a** and **6g**). Indeed, most of the observed variants were C→A (**Fig. 6b**). Consequently, S4 received the highest activity estimate by QP (93%).

**Figure 6:**
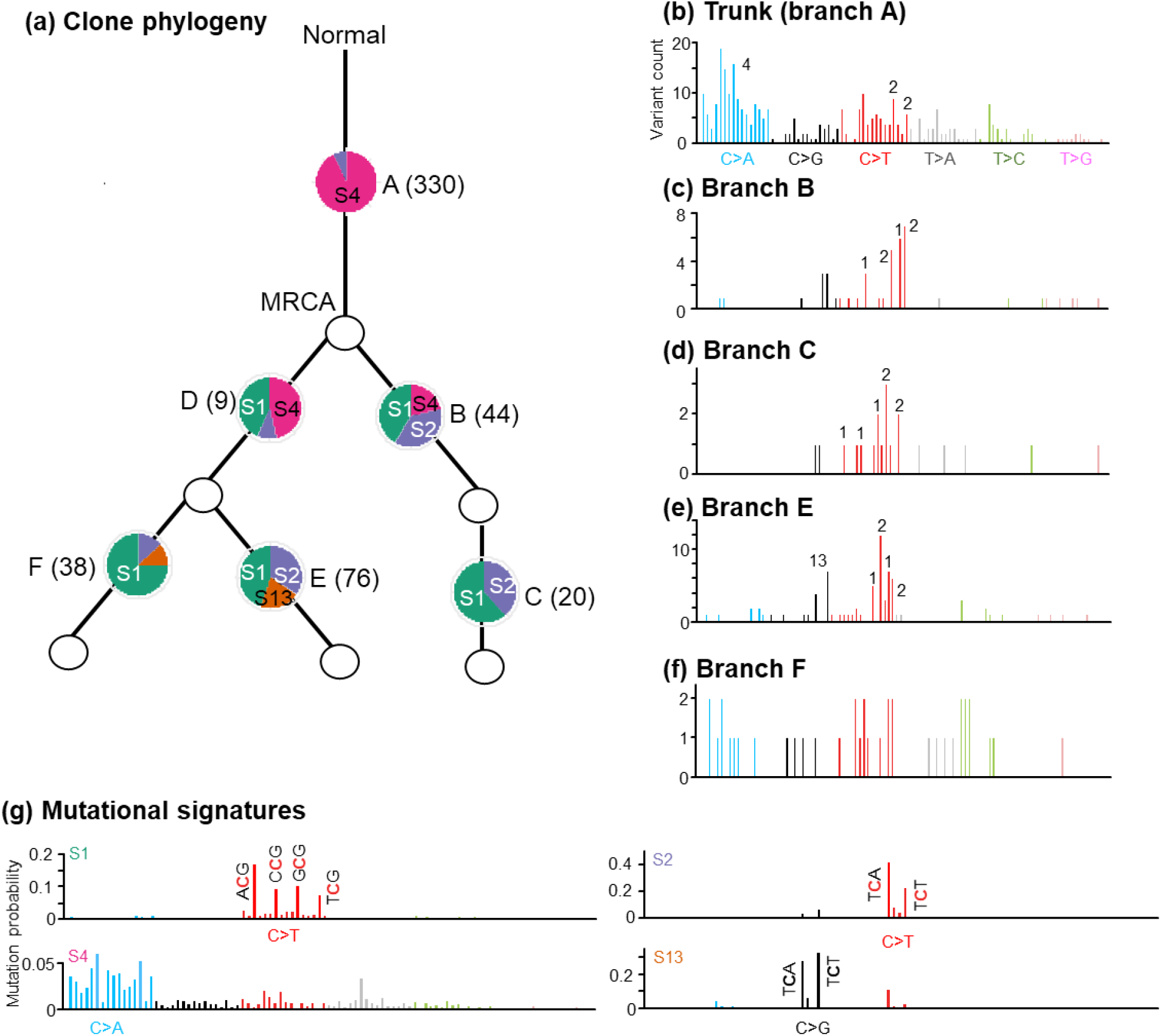
*PhyloSignare* (PS_QP_) inferences on CRUK0025 patient data. (**a**) Clone phylogeny and the mutational signatures identified for different branches (A – F). The number in the parentheses is the variant count for each branch, and a pie-chart shows the relative activities of mutational signatures. The most recent common ancestor (MRCA) of all observed clones is marked. (**b–f**) Distribution of variants observed at each branch. The numbers on top of the vertical bars correspond to variant types that were important for COSMIC signatures detected. (**g**) Distribution of variants for four COSMIC signatures that were detected for this phylogeny (S1, S2, S4, and S13).

COSMIC signature S2 was also active in the trunk, associated with the AID/APOBEC family of cytidine deaminases (Alexandrov et al., 2015; Alexandrov et al., 2020). The activity of S2 was 13 times lower than S4 in the trunk but much higher than S4 in the rest of the branches in the clone phylogeny (**Fig. 6a**). In fact, the activity of S4 was lower in the direct descendants of the MRCA, and it became too small to be detected in the tip branches C, E, and F. Therefore, the mutational processes giving rise to S4 seem to not operate later in tumor evolution (**Fig. 6a**). Another AID/APOBEC mutational signature, S13, was detected only in tip branches E and F, suggesting that it became active more recently. In comparison, the contribution of S1, age-related mutational signature, was high in all the branches in the clone phylogeny except in the trunk (**Fig. 6a**). Therefore, the evolutionary dynamic of mutational processes during clonal evolution in this patient is revealed due to the ability of *PhyloSignare* to infer branch-specific signatures even though branches are relatively short.

The evolutionary dynamics of mutational patterns for patient CRUK0025 were recapitulated in data analysis from 60 additional patients. S4 had the highest activity in the trunk of clone phylogenies of more than 72% of the patients (=44/61). Often, S4 activity declined over time, such that it became low in tips compared to the trunk (**Fig. 7a**). AID/APOBEC mutational signatures (S2 and/or S13) were also active in a vast majority of patients (>86%), with at least one of them found in the trunk branch in most patients (**Fig. 7b**). Their activity became comparable or higher than S4 in the tips. The age-related S1 signature was often gained later, or their relative activity levels became higher in tips than trunks (**Fig. 7c**). Many other COSMIC signatures associated with lung cancer (e.g., S5, S6, and S17) were detected with appreciable activity in only a small subset of patients (<30%).

**Figure 7:**
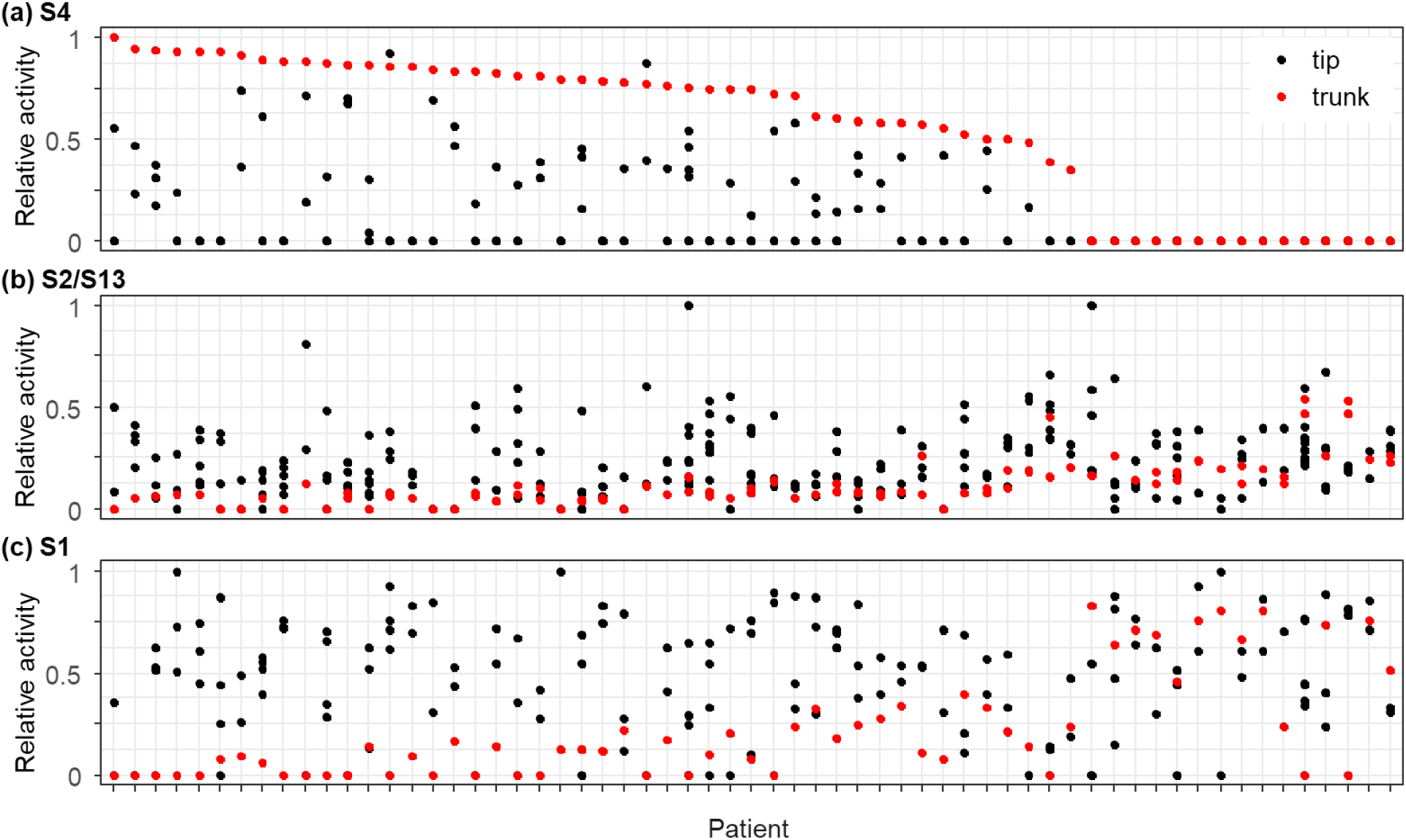
Evolutionary dynamics of mutational signatures. Relative activities of signature S4 (**a**), S2/S13 (**b**), and S1 (**c**) in the trunk (red) and tip (black) branches are shown for each patient. Patients are ordered by the relative activity of S4 in the trunk.

Within phylogenies, we conducted a more direct comparison of the presence/absence of mutational signatures between the trunk and tip branches (trunk-tip comparison) in order to quantify differences between mutational processes active in the earliest and the latest branches in patients. We constructed 162 trunk-tip comparisons. In a vast majority of pairs, there was a large difference (**Fig. 8**). The main difference was the loss/diminished activity of S4 and the gain of S1 (**Fig. 7**). That is, different sets of mutational processes were operating in the two phases of clonal evolution, which is consistent with suggestions from previous studies (de Bruin et al., 2014; Dong et al., 2018; Hao et al., 2016; Nahar et al., 2018).

**Figure 8:**
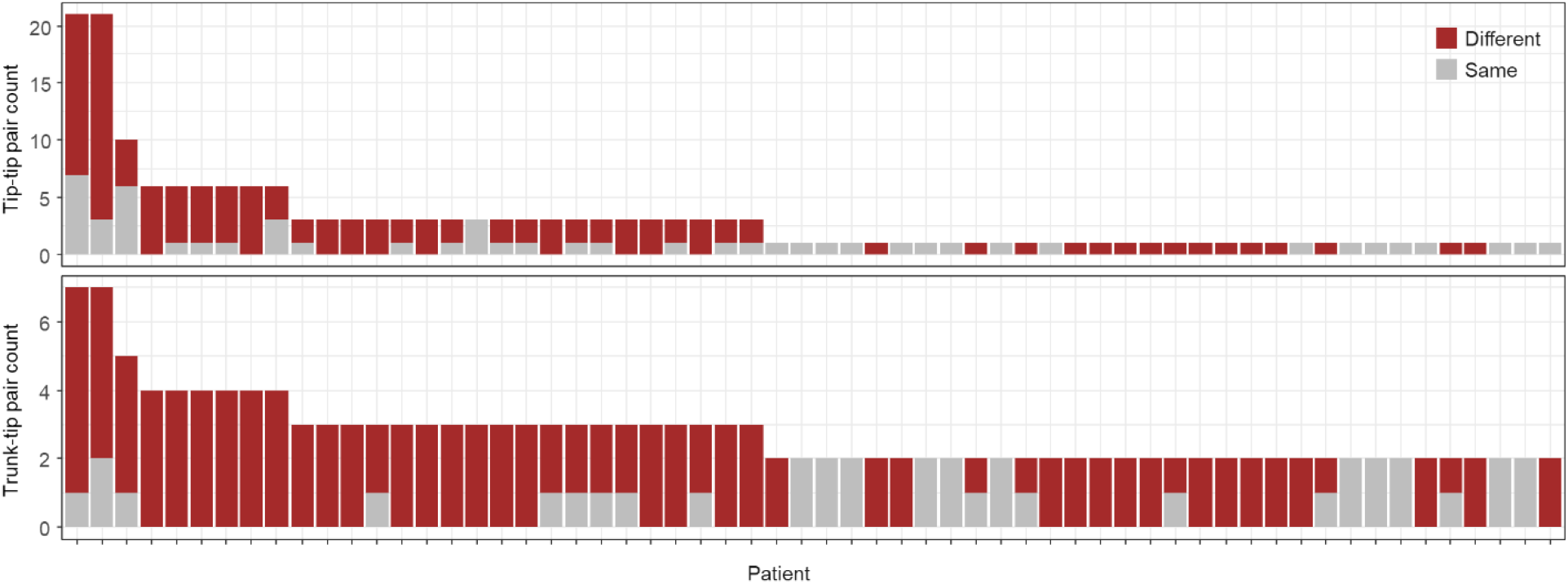
Counts of tip-tip branch pairs (top) and trunk-tip pairs (bottom) for each patient. The number of branch pairs containing different (brown) and same (gray) sets of signatures is shown. Patients are ordered based on the number of branches in their clone phylogeny.

The tip-tip comparisons provide a glimpse into the differences between the most recently diverged clonal lineages. We could conduct 176 tip-tip comparisons. We found that more than half of the clone phylogenies had at least one pair of tip-tip branches with different compositions of mutational signatures (**Fig. 8**). Common differences included the presence/absence of activities of mutational processes that give rise to signatures S1 (14/61), S2 (24/61), S4 (21/61), and S13 (22/61). Therefore, gains and losses of signatures are frequent in clone evolution, resulting in heterogeneity of mutational processes over time and space. Evolutionary patterns of signature activities varied among patients and signature types.

## DISCUSSION

Identifying lineage-specific mutational signatures has been challenging because the number of variants needed to make a reliable inference has been rather large. One way to address this problem is to conduct whole-genome sequencing (WGS) to collect hundreds of variants for each branch in the clone phylogeny (Leong et al., 2019; Yates et al., 2017). However, there may not be enough variants per branch even in genome-scale investigations if new clones frequently arise, resulting in short branch lengths or somatic evolution has been occurring for a short period of time or with a slow rate. Also, currently, exome sequencing is often used in research investigations, which means that the number of variants mapped to individual branches may not be large enough for existing methods. This means that *PhyloSignare* will likely be useful in most future investigations. We have found that it is possible to improve the quality of mutational signature identification for individual branches of clone phylogenies.

The use of *PhyloSignare* provided new insights into clonal evolution using the existing data compared to those derived previously for the same datasets (Jamal-Hanjani et al., 2017). Jamal-Hanjani et al. (2017) used a standard signature activity estimation method, followed by manual curation, to identify at most one mutational signature for every branch. For example, in the patient CRU0025, they reported the presence of S4 in the trunk (A) and S2/S13 its two descendants (B and E; see **Fig. 6**). No mutational signatures were detected for branches C, D, and F. Using *PhyloSignare*, we found more signatures for every branch, each estimated to have significant relative activity (**Fig. 6a**). For example, S1 and S4 both have estimated relative activities similar to S2 in branch B. Therefore, additional signatures were made detectable for branches by using *PhyloSignare*. With high precision, *PhyloSignare* makes it possible to detect changing dynamics of mutational processes over time in a patient. We find that mutational signature patterns across patients show convergence towards a loss of smoking-related signatures, consistent with previous lung cancer evolution reports (de Bruin et al., 2014). We also found a convergent tendency to gain AID/APOBEC signatures in MRCA’s descendants, which suggests that mutational processes begin shifting while the tumor cells diversify from the MRCA over time. There is also a tendency for mutational signatures to diverge among closely related lineages (e.g., tip-tip pairs), suggesting regional and/or temporal differences in the mutational and selective pressures within tumors.

We did not always detect S1, associated with aging, in the trunk, but S1 was otherwise found in a majority of branches in the phylogeny. S1’s ubiquity is reasonable because the mutational processes due to aging should be present throughout. But, its detection in the presence of S4 seems to be difficult because of the much stronger activity of S4 that likely overwhelms S1’s signal because C→T mutations are common to both S1 and S4. In small sample sizes, distinguishing between S1, S2, and S6 is also difficult because they involve C→T mutations. So, the absence of a lung-cancer-related S6 signature in the Jamal-Hanjani et al. (2017)’s dataset could be due to the detection problem. Another lung-related signature, S5, was also not often detected because it is a flat signature (i.e., many different types of mutations occur with a similar probability), whose detection is notoriously difficult even with large variant collections (Maura et al., 2019). Therefore, the absence of some expected lung-cancer signatures does not mean that those mutational processes are inactive, and statistical methods need to be improved to detect them.

In conclusion, *PhyloSignare* improves the accuracy of mutational signature detected using standard methods for smaller variant collections. Its application reveals dynamics of mutational signatures at a higher phylogenetic resolution in this cancer, enabling the detection of mutational activity over time and among closely-related lineages.

## MATERIAL AND METHODS

### PhyloSignare approach

We describe this approach using an example clone phylogeny (**Fig. 4a**) consisting of five branches onto which 20–100 variants were mapped (a general flowchart is provided in **Fig. 9**). First, we identify candidate signatures for each branch by applying a user-selected mutational signature detection method, e.g., quadratic programming (QP) technique (Huang et al., 2018), DeconstructSigs (dSig) (Rosenthal et al., 2016), and MutationalPatterns (MutPat) (Blokzijl et al., 2018). For example, to obtain candidate signatures for branch B, one would apply the selected mutational signature detection method (QP used here) to (1) variants within branch B, (2) a pooled collection of variants from branches B and C (B’s sibling branch), (3) a pooled collection of variants from B and A (B’s direct ancestral branch), and (4) pooled collection of variants from B, A, and C. The objective of pooling information with neighboring branches is to increase the number of variants that enhance existing methods’ statistical power to detect mutational signatures with low activity. By using QP, we obtain the activity of all COSMIC signatures (v2) in these collections. Mutational signatures with estimated activity greater than 0.01 in at least one collection were included to assemble a set of candidate signatures for branch B. We selected this 0.01 cutoff value because almost half of the incorrect signatures that QP detected had <0.01 estimated relative activities in our simulation study.

**Figure 9.**
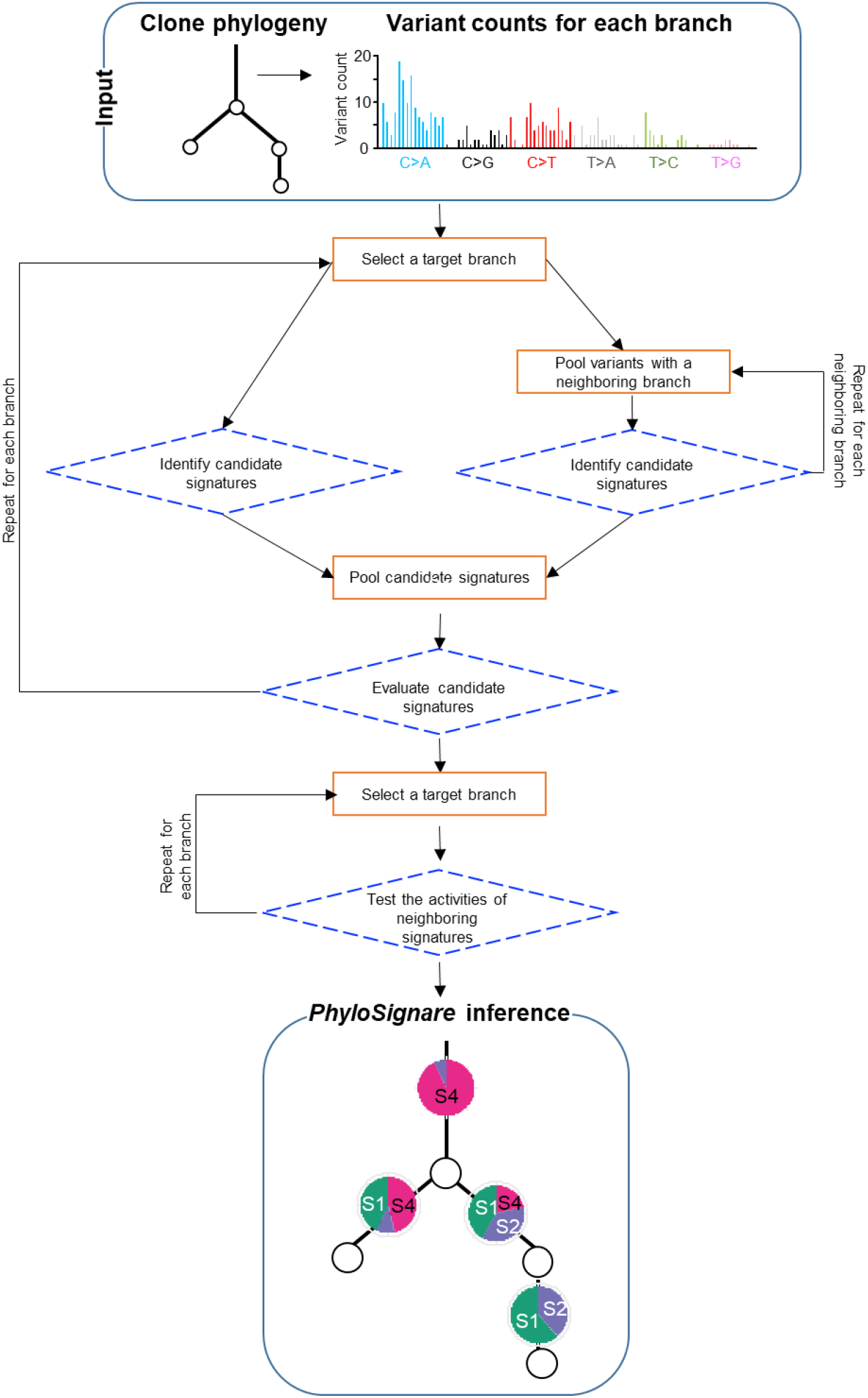
Overview of *PhyloSignare* approach. Our approach uses a clone phylogeny in which all variants are mapped along branches. *PhyloSignare* pools mutations with adjacent branches and collect candidate signatures for each branch. We use iS statistic (see text) to evaluate the presence of each candidate signature. Last, we test if signatures from neighboring branches are active at a branch. Signatures will be detected for each branch.

We next test the significance of the predicted signature activities. For every candidate signature (*S*), we compute a simple importance score (iS), iS = (*f_s–_* – *f*)*/f*, where, 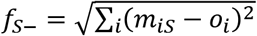. The *m_is_* is estimated for every variant after signature *S* is excluded, which is a product of the mutational signature matrices specified, estimated relative activities, and the total mutation count. The other term is, 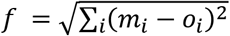, where *m_i_* is an estimated mutation count when signature *S* is included. Basically, iS is expected to be close to zero if a given signature *S* is spurious, i.e., such signatures are unlikely to contribute significantly to the fit of the observed data; we retain signatures with iS > 0.02. In the final step, the final candidate list of signatures for a branch is built by pooling signatures inferred for that branch along with the signatures of its immediate relatives. Then, we estimate the relative activities of these signatures using QP. This step is meant to minimize spurious gain and loss of signatures caused by small sample size. The *PhyloSignare* software is available at https://github.com/SayakaMiura/PhyloSignare.

By the way, in the above, we assumed that the clone phylogeny is known. In empirical data analysis, one needs to generate it using available computational tools for bulk and single-cell sequencing methods; see reviews in the accuracy of methods (Miura et al., 2018a; Miura et al., 2018b; Miura et al., 2020). The errors in the collection of variants for each branch will lead to false-negative detection of signatures due to diluted signals caused by incorrect variants and correct variants that are not assigned to a branch. Therefore, we encourage users to scrutinize the quality of inferred clone phylogenies before applying *PhyloSignare*.

### Collection and analysis of simulated datasets

We obtained 180 simulated datasets from the website https://github.com/elkebir-group/PhySigs (Christensen et al., 2020). Each clone phylogeny (containing five or seven branches) can be partitioned into up to three subtrees, each with an identical set of mutational signatures and relative activities. Each branch of these clone phylogenies had from 2 to 205 mutations. COSMIC v2 signatures were randomly sampled to generate these datasets, and relative exposures at each branch were determined by drawing from a symmetric Dirichlet distribution. Observed mutation counts at each branch were generated by introducing Gaussian noise with a mean of zero and standard deviation of 0.1, 0.2, or 0.3. Some of the simulated signature activities were small (<10%). Since the detection of these signatures is impossible, especially when the number of mutations is small, we did not consider them as a failure of the detection component of a method when they were not detected.

We applied *PhyloSignare* to these simulated datasets by providing correct clone phylogenies and v2 COSMIC signatures obtained from https://cancer.sanger.ac.uk/cosmic/signatures. For each branch mutation count, we also performed QP (Huang et al., 2018), dSig (Rosenthal et al., 2016), and MutPat (Blokzijl et al., 2018) by providing v2 COSMIC signatures. Here, signatures that were estimated with <0.001 relative frequencies were considered to be absent. dSig was performed by using the option to discard inferred signatures with <0.001 relative frequencies. We excluded branches with <20 variants from the accuracy evaluation because signature detection is impossible for any method.

Our simulation study excluded methods that were not designed to estimate the relative activities of signatures for each branch. For example, we did not include PhySigs (Christensen et al., 2020) because PhySigs is designed to classify branches based on similarities of variant counts, and relative activities of signatures are produced as a byproduct. In any case, the true positive rate of PhySigs was much lower than *PhyloSignare* (precision equal to 71%), and the overall accuracy of *PhyloSignare* was better (F1 equal to 91% and 83% for *PhyloSignare* and PhySigs, respectively); PhySigs inferences were obtained from https://github.com/elkebir-group/PhySigs.

### Collection and analysis of empirical datasets

We obtained 100 non-small cell lung cancer (NSCLC) data from the TRACERx Lung Cancer study (Jamal-Hanjani et al., 2017). We collected only invasive adenocarcinoma and squamous cell carcinoma samples (61 and 32 samples, respectively) because the number of the other cancer types was very small. These datasets contained inferred clone phylogenies with all observed mutations mapped along branches. We selected the primary phylogenies when more than one phylogenies were reported. We then excluded datasets when the total number of variants was less than 100 or when a clone phylogeny did not have at least two tip branches. After these filtering processes, we obtained clone phylogenies from 61 patients.

We classified each observed mutation into the 96 trinucleotide mutation patterns and generated branch-specific mutation counts used as input information for *PhyloSignare*. When a mutation count for a branch was < 20, we pooled them with its neighboring branch because it was impossible to identify mutational signatures on data with a number of mutations too small (red branches in **Fig. 2**). The input files for *PhyloSignare* are deposited at https://github.com/SayakaMiura/PhyloSignare. To perform the *PhyloSignare* analysis, we used COSMIC v2 signatures known in lung adenocarcinoma (S1, S2, S4, S5, S6, S13, and S17) and squamous signatures (S1, S2, S4, S5, S13). Accordingly, we provided each set of known signatures in the analysis based on the given dataset’s cancer type. We used QP to estimate relative activities in all our data analyses.

## ACKNOWLEDGMENTS

We thank Drs. Marcos Caraballo and Antonia Chroni for comments and editorial support. We also thank Vivian Aly and Sudip Sharma for technical support. This research was supported by the National Institutes of Health to S.K (LM012487) and S.M. (LM012758).

